# Developmental Characterization of *Zswim5* Expression in the Progenitor Domains and Tangential Migration Pathways of Cortical Interneurons in the Mouse Forebrain

**DOI:** 10.1101/728097

**Authors:** Chuan-Chie Chang, Hsiao-Ying Kuo, Shih-Yun Chen, Kuan-Ming Lu, Weng Lam Fong, Hsiao-Lin Wu, Tetsuichiro Saito, Fu-Chin Liu

## Abstract

GABAergic interneurons play an essential role in modulating cortical networks. The progenitor domains of cortical interneurons are localized in developing ventral forebrain, including the medial ganglionic eminence (MGE), caudal ganglionic eminence (CGE), preoptic area (POA) and preoptic hypothalamic border domain (POH). Here, we characterized the expression pattern of *Zswim5*, an MGE-enriched gene in the mouse forebrain. At E11.5 to E13.5, prominent *Zswim5* expression was detected in the subventricular zone (SVZ) of MGE, CGE, POA and POH of ventral telencephalon in which progenitors of cortical interneurons resided. At E15.5 and E17.5, *Zswim5* remained detectable in the SVZ of pallidal primordium (MGE). *Zswim5* mRNA was markedly decreased after birth and was absent in the adult forebrain. Interestingly, *Zswim5* expression pattern resembled the tangential migration pathways of cortical interneurons. *Zswim5*-positive cells in the MGE appeared to migrate from the MGE through the SVZ of LGE to overlying neocortex. Indeed, *Zswim5* was co-localized with Nkx2.1 and Lhx6, markers of progenitos and migratory cortical interneurons. Double labeling showed that Mash1/Ascl1-positive cells did not express *Zswim5*. *Zswim5* expressing cells showed none or at most low levels of Ki67 but co-expressed Tuj1 in the SVZ of MGE. These results suggest that *Zswim5* is immediately upregulated as progenitors exiting cell cycle to become postmitotic. Given that recent studies have elucidated that the cell fate of cortical interneurons is determined shortly after postmitotic, the timing of *Zswim5* expression in early postmitotic cortical interneurons suggests a potential role of *Zswim5* in regulation of neurogenesis and tangential migration of cortical interneurons.

## INTRODUCTION

Propagation of neuronal information in neural networks is modulated by the balance between excitatory and inhibitory signals. Abnormalities in excitatory/inhibitory (E/I) balance of synaptic activity have been well documented in neurodevelopmental disorders such as autism and schizophrenia (Ramamoorthi and Lin, 2011; Nelson and Valakh, 2015; Canitano and Pallagrosi, 2017; Sohal and Rubenstein, 2019). A large diversity of cortical interneurons with distinct morphology, connectivity, and physiological activity serves to regulate synaptic E/I balance of cortical projection neurons. Understanding the developmental roots of GABAergic interneurons should provide important information to the pathophysiology of the diseases associated with E/I imbalance.

GABAergic interneurons in the pallium of telencephalon are well known to be developmentally derived from the subpallium of the telencephalon (Marin and Rubenstein, 2001; Marín, 2015; Bandler et al., 2017; Hu et al., 2017; Lim et al., 2018). In the ventral part of developing mammalian telencephalon (subpallium), there are three structural elevations, the lateral ganglionic eminence (LGE), medial ganglionic eminence (MGE) and the caudal ganglionic eminence (CGE). These three ganglionic eminences consist of heterogeneous progenitor populations that give rise to neurons in the striatum, globus pallidus, amygdala, and other basal forebrain regions. Importantly, the MGE, CGE, and LGE also give rise to GABAergic interneurons in the cerebral cortex, hippocampus and striatum through long-range tangential migration from the subpallium to the pallium (Marin and Rubenstein, 2001; Miyoshi et al., 2013; Marín, 2015; Bandler et al., 2017; Hu et al., 2017; Lim et al., 2018).

Previous studies have documented that most cortical GABAergic interneurons are born in the MGE, CGE, preoptic area (POA) and preoptic hypothalamic border domain (POH) (Wonders and Anderson, 2006; Miyoshi et al., 2013; Marín, 2015; Bandler et al., 2017; Hu et al., 2017). The MGE comprises major pools of cortical GABAergic interneurons that are regulated by the MGE-enriched Nkx2.1 gene (Sussel et al., 1999; Sousa and Fishell, 2010). Somatostatin (SST)- and parvalbumin (PV)-positive interneurons are mainly generated from the dorsal and ventral parts of MGE, respectively, and SST-positive interneurons are generated earlier than PV-positive interneurons (Miyoshi et al., 2007; Inan et al., 2012). The ventral-most MGE region produces striatal interneurons (Wonders and Anderson, 2006; Flames et al., 2007). Vasointestinal peptide (VIP)-, reelin (RELN)-, calretinin-, neuropeptide Y (NPY)-, and 5HT3a receptor-positive cortical interneurons are derived from the CGE (Xu et al., 2004; Butt et al., 2005; Lee et al., 2010; Miyoshi et al., 2010; Rudy et al., 2011; Marín, 2015; Lim et al., 2018). Small populations of cortical interneurons and olfactory bulb interneurons are generated from the LGE (Stenman et al., 2003; Xu et al., 2004). NPY-, PV- or SST-positive cortical interneurons that are generated at the earliest time are mainly derived from the POA in which interneuron progenitors also express Nkx2.1 (Flames et al., 2007; Gelman et al., 2011). Unlike the POA, the POH shares molecular similarities with the CGE and generates 5HT3a receptor-positive cortical interneurons, including RELN-positive neurogliaform cells and NPY-positive multipolar cells (Lim et al., 2018; Niquille et al., 2018). After being generated in the progenitor domains, postmitotic cortical interneurons tangentially migrate from the subcortical regions through the migratory streams to the developing cortex.

In the present study, we investigated *Zswim5*, an MGE-enriched gene that has not been well characterized in the developing forebrain. The Zswim family is a class of genes containing a SWIM domain that is structurally similar to the zinc-finger motif. The SWIM domain is characterized by a CxCxnCxH motif of predicted zinc chelating residues, and is conserved in both prokaryotic and eukaryotic proteins (Makarova et al., 2002). In the CxCx_n_CxH motif, the “C” stands for cysteines, and the “H” stands for histidines. Both “C” and “H” are predicted to serve as chelating metal residues in this motif. The “n” in CxCx_n_CxH typically varies between 6 to 16 residues, whereas some proteins may have up to 25 residues. This conserved pattern was first identified when aligning the family of bacterial SWI2/SNF2 ATPases of the helicase superfamily II (Pazin and Kadonaga, 1997). More than 100 protein sequences containing the CxCx_n_CxH signatures were identified through the PSI–BLAST searches (Altschul et al., 1997). Among them, three experimentally characterized protein sequences were also identified, including the plant MuDR transposases (Hershberger et al., 1995; Benito and Walbot, 1997), the FAR1 family of plant nuclear proteins involved in phytochrome signal transduction (Hudson et al., 1999) and the vertebrate MEK kinase-1 (MEKK-1) (Hagemann and Blank, 2001). Therefore, the SWIM domain is named after <u>SWI</u>2/SNF2 and <u>M</u>uDR transposases. Although only a small core of the SWIM domain shows sequence conservation, it appears to have a ββα structure, suggesting that it might adopt a fold similar to that of the classic C2H2 Zinc-finger (Makarova et al., 2002). Functionally, it appears to be a versatile domain predicted to interact with either DNA or proteins in different contexts. Further experimental studies on the SWIM domain will reveal how this common structural scaffold is used in apparently different processes, such as MuDR transposition in plants and MEK kinase signaling in animals (Makarova et al., 2002).

*Zswim5* (MGI: 1921714; transcript: NM_001029912; polypeptide: Q80TC6) is located on chromosome 4, and it is also known as mKIAA1511 in the mouse KIAA cDNA database (see below). *ZSWIM5* protein comprises 1188 a.a., and the SWIM domain is speculated to locate at 222-259 a.a. (YKVAISFDRCKITSVS**<u>C</u>**G**<u>C</u>**GNKDIFY**<u>C</u>**A**<u>H</u>**VVALSLYRI, 38 a.a.). Previous studies suggest that *Zswim5* may be involved in neural crest formation and high-grade human glioma (Meyer, 2014; Wong et al., 2016). However, the biochemical property and physiological function of *Zswim5* protein are largely unknown.

Previous microarray analysis has found that the expression level of *Zswim5* in the MGE is 2.8-fold higher than that in the LGE (Tucker et al., 2008). Recent high-throughput single-cell RNA sequencing studies have also identified *Zswim5* as a maker of progenitors of cortical interneurons (Mayer et al., 2018; Mi et al., 2018), but the spatial and temporal expression pattern of *Zswim5* remains elusive. Here, we performed *in situ* hybridization using digoxigenin-labeled and ^35^S-UTP-labeled riboprobes to delineate the spatiotemporal expression pattern of *Zswim5* mRNA in the mouse forebrain during development. We found that *Zswim5* was expressed in differentiating progenitors of cortical GABAergic interneurons that were undergoing tangential migration, suggesting that a potential regulation of the development of cortical interneurons by *Zswim5*.

## MATERIALS AND METHODS

### Animals

The mice used in this study were kept in the animal center of National Yang-Ming University and followed the protocols of animal use approved by the Institutional Animal Care and Use Committee. Efforts were made to minimize the suffering and the number of animals used. To define the stages of mice, noon on the day with a vaginal plug was considered as embryonic day 0.5 (E0.5) and the day of birth as postnatal day 0 (P0). For this study, Imprinting Control Region (ICR) mice brains at different embryonic (E11.5, E12.5, E13.5, E15.5, and E17.5) or postnatal (P0, P7, P14, and adult) stages were harvested according to the following protocol. Time-pregnancy ICR mice were first deeply anesthetized with 0.05∼0.08 ml pentobarbital (i.p. injection), and the embryos were release from its V-shaped uterus with forceps and scissors. The head of E11.5, E12.5, and E13.5 ICR embryos were directly cut off from the body and immediately fixed with 4% paraformaldehyde in 1X Phosphate Buffered Saline (PBS, pH 7.4) and gently shake at 4°C overnight. For E15.5 and E17.5, brains were dissected out from the head and fix with 4% PFA/1X PBS overnight at 4°C. As for postnatal stages, including P0, P7, P14, and adult (around 2 months), brains were perfused with 0.9% saline and 4% PFA/1X PBS. Following post-fixation, brains were cryoprotected in 30% sucrose in 1X PBS at 4°C for 2 to 3 overnights. All brains were individually wrapped in aluminum foil and instantly frozen by dry ice and then stored at −70°C for further processing.

### Cryo-sectioning of brain tissue

Frozen brains were taken out from −70°C and put into a rubber-made container filed with Cryo-Gel (Instrumedics). The container was then put on dry-ice to freeze the brain sample as quickly as possible to prevent degradation of RNA or proteins. Using rapid sectioning cryostat (Leica CM1900), 20 μm or 25 μm thick cryosections were collected and pasted on silane-coated slides (DAKO). After sectioning, the slides were air-dried for a few hours, and then collected in a clean slide box and stored in −20°C for further usage.

### The template plasmids for riboprobe synthesis

To obtain the specific expression pattern of *Zswim5* mRNA in the mouse brain during development, the specific partial sequence of *Zswim5* was selected to synthesize probes *in vitro*, which theoretically would minimized the chance of recognizing other *Zswim* family members. The *Zswim5, Nkx2.1* and *Lhx6* plasmids for riboprobe synthesis are listed in Table 1.

**Table 1.** Summary of the cRNA probes for in situ hybridization

| Gene | Vector | Clone name | Probe size | Anti-sense |  | Sense |  | Hybridization temperature |
| --- | --- | --- | --- | --- | --- | --- | --- | --- |
| Lhx6 | pGEM-T easy | pGEM-Lhx6 | 350 bp | NcoI | Sp6 | – | – | 65°C |
| Nkx2.1 | pGEM-T easy | pGEM-Nkx2.1 | 328 bp | NcoI | Sp6 | – | – | 65°C |
| Zswim5 | pGEM-T easy | pGEM-Zswim5-1807 | 1807 bp | NcoI | Sp6 | Sall | T7 | 65°C |
|  | pBC SK(+) | pBCSK(+)-Zswim5 | Full length | XhoI | T3 | NotI | T7 | 65°C |

### Subcloning of *Zswim5* partial sequence into the pGEM-T easy vector

The *Zswim5* full-length cDNA clone constructed in pBC SK+ vector (mKIAA1511) was kindly provided by Dr. Hisashi Koga and Takahiro Nagase at the Kazusa DNA Research Institute in Japan (Okazaki et al., 2003). The *Zswim5* cDNA fragment (5,401 bps) was inserted between XhoI and NotI site of the Chloramphenicol-resistant pBC SK+ vector (3,400 bps). The targeted partial *Zswim5* fragment was first amplified by PCR using forward (Z5-3’UTR-5’) and reverse (Z5-3’UTR-3’) primers: 5’-ctggg caaga atgaa ctggc-3’ and 5’’-aatac cagcc tcagc ctccg-3’, respectively. With total volume of 50 μl, every 10 μl PCR reaction contains 4.7 μl 3dH_2_O; 2 μl 5.5M betaine; 0.5 μl 5’ primer (10 mM), 0.5 μl 3’ primer (10 mM), 0.2 μl dNTP (25 mM), 1 μl 10X PCR buffer, 1 μl plasmid and 0.1 μl Taq polymerase (Geneaid). After thorough homogenization, a total volume of 50 μl was equally divided into 5 tubes to amplify the target fragment under the following PCR condition: 94°C for 3 min, 30 cycles of denaturing step (94°C for 30 seconds), primer annealing step (60°C for 30 seconds), and amplifying step (72°C for 1 min), 72°C for 5 min, and finally stored at 4°C. An 1807 bps-long fragment was amplified and further eluted out with elution Kit (Geneaid). After confirming the size and the molecular weight of the PCR product with 1Kb marker, the target *Zswim5* partial fragment was ligated with pGEM-T easy vector by T4 ligase (NEB). With the vector: insert ration equals to 1:3 rule, total volume of 15 μl PCR mixture was incubated at 16°C overnight. The ligation product was then transformed in the competent cell DH5α (Yeastern Biotech) and screened for clones that were resistance to Ampicillin. Clones with correct insertions were selected by PCR with forward (Z5-3’UTR-5’) and reverse (SP6) primers: 5’-ctggg caaga atgaa ctggc-3’ and 5’-attta ggtga cacta tag-3’, respectively. Following PCR condition: 94°C for 3 min, 30 cycles of denaturing step (94°C for 30 seconds), primer annealing step (50°C for 30 seconds), and amplifying step (72°C for 2 min), 72°C for 2 min, and finally stored at 4°C. The positive clones were further checked with restriction enzymes NotI (GC^▾^GGCCGC, NEB) and PstI (CTGCA^▾^G, NEB), generating fragments of 1,851 bps; 2,977 bps and 1,246 bps; 3,583 bps, respectively.

### Synthesis of non-radioactive RNA probes

The non-radioactive RNA probes were synthesized by *in vitro* transcription with digoxigenin (dig) or fluorescein (FITC) RNA labeling mix (Roche). In brief, respective linearized template plasmid (1 μg) is mixed with 4 μl 5X transcription buffer (Promega), 2 μl 0.1M DTT (Promega), 2 μl Dig- or FITC-labeling mix (Roche), 1 μl RNasin (Promega), 1.5 μl respective RNA polymerase (T3, T7, SP6, Promega) and DEPC H_2_O in a total volume of 20 μl at 37°C for 2 hr. The DNA template is digested with the following DNase RQ1 treatment at 37°C for 30 min. The polymerase reaction is stopped by adding 5 μl 0.2M EDTA (pH 8.0) and stayed on the ice for 5 min. After adding 30 μl STE buffer (10 mM Tris-HCl, pH 8.0; 1 mM EDTA, pH8.0; 0.1 M NaCl) and 3 μl 1M DTT, the labeling product is further purified with G-50 mini Quick Spin Columns (Roche). The final qualities of probes were evaluated through the products sampled before DNase RQ1 (Promega) treatment and after the column purification. For better performance, the probes were aliquot in a small volume and stored at −70°C before hybridization.

### Non-radioactive *in situ* hybridization

Slides were air-dried at room temperature for 10 min and vacuumed in the desiccators for at least one hour to ensure no water remained on the sections. For embryonic stages, sections were washed in 1X PBS for 5 min and treated with 0.1% Triton X-100 in 1X PBS for 5 min to remove the lipid. For P0 to adult stages, sections were post-fixed in 4% PFA in 1X PBS for 30 min on ice and then treated with 0.3% Triton X-100 in 1X PBS for 15 min. After washing in 1X PBS, all sections were incubated in 0.2N HCl in DEPC H_2_O for 20 min. Crucially, sections at different stages were treated with proteinase K (PK, 10 μg/ml, MDBio, Inc.) in 1X PBS at 37°C for 2 to 5 min for protein removal. Following washing in 1X PBS, sections were fixed with 4% PFA/1X PBS for 5 min and incubated twice in glycine (2 μg/ml) in 1X PBS for 15 min. Sections were then prehybridized with 50% deionized formamide (Sigma) in 2X standard saline citrate (SSC, 30 mM sodium citrate, 300 mM NaCl, pH 7.0) at 65°C for 90 min in a humid box. Dilute the probes with ratios ranging from 1:250 to 1:1000 in the hybridization solution I and II (10% dextran sulfate; 50% formamide; 1 mM EDTA, pH 8.0; 0.01M Tris, pH 8.0; 0.3M NaCl, 1X Denhardt’s solution, 500 μg/ml yeast tRNA and 10 mM DTT) and denature the probe at 90°C for 10 min. Each slide is added with 200 μl hybridization solution containing respective probe and covered with a coverslip and sealed with rubber cement. After 16 hr of hybridization at 65°C (D1R and D2R at 60°C), sections were washed in 5X SSC for 5 min and then incubated in 50% formamide (Sigma) in 2X SSC for 1 hour. Then sections were incubated in 10 mM Tris-HCl (pH 8.0) and 500 mM NaCl for 10 min before and after treated with RNase A (20 μg/ml) at 37°C for 30 min. The sections were washed sequentially in 2X SSC, and 0.2X SSC twice for 20 min at 65°C. After washing with TNT buffer (100 mM Tris pH 7.5, 150 mM NaCl) for 10 min, sections were blocked with 2% blocking reagent, 20% sheep serum in the TNT buffer for 60 min. The Dig-labeled probes hybridized with target transcripts were recognized by alkaline phosphatase (AP)-conjugated sheep anti-digoxigenin antibody (1:1000, 11093274910, Roche, Basel, Switzerland, www.roche.com, RRID: AB_514497) for 90 min. After washing sections in TNT buffer and buffer 3 (100 mM Tris-HCl pH 9.5, 100mM NaCl), signals were detected by colorimetric reaction using Nitro blue tetrazolium chloride (NBT, Roche) and 5-Bromo-4-chloro-3-indolyl phosphate (BCIP, Roche) in buffer 3 as the substrates.

For fluorescent system, all TNT buffer is added with 0.1% Tween-20. After treated with 0.1% H_2_O_2_ in TNT, which removed the endogenous peroxidase, sections were blocked with 2% blocking reagent, 20% sheep serum in the TNT for 60 min. Antibodies used to detect the probes were horseradish peroxidase (HRP)–conjugated sheep anti-digoxigenin (1:100, 11207733910, Roche, Basel, Switzerland, www.roche.com, RRID: AB_514500) or HRP-conjugated sheep anti-FITC antibody (1:100, 11426346910, Roche, Basel, Switzerland, www.roche.com, RRID: AB_840257). After incubating overnight in the antibody, the signals were further detected with the Tyramide Signal Amplification System (TSA, PerkinElmer) on the next day. Briefly, tyramide-Cy3 or tyramide-FITC were diluted in the 1X dilution buffer (Vector, 1:1000) and applied to the sections for 10 min. After washing with TNT buffer, the signals could be observed with conventional fluorescence microscopy (Eclipse E800M, Nikon) or confocal microscopy (TCS SP2 confocal, Leica).

For double *in situ* hybridization, the HRP activity of the first antibody used to identify the first type of transcript, usually the sheep anti-FITC antibody (1:100, 11426346910, Roche, Basel, Switzerland, www.roche.com, RRID: AB_840257), was bleached by 0.1% H_2_O_2_ in TNT for 15 min after a successful detection of fluorescent signals. The bleaching reaction was stopped by washing in TNT. Then the sections were again blocked with 2% blocking reagent, 20% sheep serum in the TNT for 60 min. Finally, the second antibody used to detect the other type of target transcripts; usually, the sheep anti-Dig antibody (1:100, 11207733910, Roche, Basel, Switzerland, www.roche.com, RRID: AB_514500) was applied to the sections for overnight incubation. Follow the same color detection method with another color using the TSA system, the double labeling of two different mRNA transcripts were distinguished and ready for further analysis.

### Radioactive *in situ* hybridization

After vacuumed in the desiccators for 60 min, the sections were fixed with 10% formaldehyde in 1X KPBS (1.5 M NaCl, 0.03 M KH_2_PO_4_, and 0.2 M K_2_HPO_4_) and followed by treatment with 10 μg/ml proteinase K (PK, MDBio, Inc.) in PK buffer containing 0.1 M Tris (pH 8.0) and 0.05 M EDTA (ph 8.0) at 37°C for 20 min. After rinsing in DEPC-H_2_O for 3 min, the sections were treated with 0.1M Triethanolamine (TEA, pH 8.0) for 3 min, followed with acetic anhydrate in 0.1M TEA for 10 min and then washed with 2X SSC buffer. Following sequential dehydration with ethanol (50%, 70%, 95%, and 100% for twice, each for 3 min), the sections were air-dried for 2 hr and further hybridized with corresponding ^35^S-labeled antisense probes. In brief, the probe was mixed with hybridization solution I and II (10% dextran sulfate; 50% formamide; 1mM EDTA, pH 8.0; 0.01M Tris, pH 8.0; 0.3M NaCl, 1X Denhardt’s solution, 500 μg/ml yeast tRNA and 10mM DTT) in a ratio of 10^7^cpm ^35^S-UTP-cRNA per ml solution. After denaturing the probe at 65°C for 5 min, the hybridization mix was applied to each slide and hybridized at 58°C for 16 hr. On the next day, after washing with 4X SSC for 7 min four times, the sections were treated with 10 μg/ml RNase A in buffer containing 10 mM Tris (pH 8.0), 0.5M NaCl, and 1mM EDTA (pH 8.0) at 37°C for 30 min. Then, the sections were subsequently washed in 2X SSC for twice, 1X SSC and 0.5X SSC for once, each for 5 min. The slides were then incubated in 0.1X SSC at 50°C for 30 min and 0.1X SSC at room temperature for 5 min. All SSC solutions were added with 1mM DTT. After dehydrated with EtOH (50%, 70%, 95%, and 100% twice) and vacuumed in the desiccators for at least 2 hr, the sections were exposed to X-ray file to visualize the ^35^S isotope signals by autoradiography.

### Immunohistochemistry

For single-labeling immunohistochemistry without antigen retrieval, cryo-sections were first treated with 0.1% sodium azide in 0.1M PB for at least 30 min. After washing with 0.1M PBS, sections were treated with 0.2% Triton-X100 in 0.1M PBS for 10 min to permeabilize the cell membrane, which allows the antibody to penetrate and reach for its target antigen within the cell. Treating with 3% H_2_O_2_, 10% methanol, and 0.2% Triton-X100 in 0.1M PBS for 5 min, endogenous peroxidase within the cell is bleached out to prevent a non-specific color reaction in the signal detecting process. After washing with 0.1M PBS, sections were blocked with 3% normal goat serum (NGS) in 0.1M PBS for 1 hour to prevent non-specific binding of the antibody. Different primary antibodies diluted with proper titrations in 0.1M PBS containing 1% NGS, 0.2% triton-X100 and 0.1% sodium azide were then applied to the cryo-sections and further incubated at room temperature for overnight. The information of all primary antibodies containing rabbit anti-Ki67 (1:200, NCL-Ki67p, Leica Biosystems, Illinois, United States, www.leicabiosystems.com, RRID: AB_442102), rabbit anti-Lhx6 (1:50, ab22885, Abcam, Cambridge, United Kingdom, www.abcam.com, RRID: AB_447345), rabbit anti-Lhx8 (1:2000, ab41519, Abcam, Cambridge, United Kingdom, www.abcam.com, RRID: AB_943992), mouse anti-Tuj1 (1:4000, G7121, Promega, Wisconsin, United States, www.promega.com, RRID:AB_430874) and mouse anti-Mash1 [1:100, a gift of Prof. D. J. Anderson in California Institute of Technology, Pasadena, United States. This antibody has been previously characterized and published (Lo et al., 1991)]. Primary antibodies were washed out with 0.1M PBS on the following day. After incubation with secondary antibodies conjugated with fluorescent materials, such as FITC conjugated goat-anti-rabbit (1:250, 111-095-003, Jackson ImmunoResearch, Pennsylvania, United States, www.jacksonimmuno.com, RRID: AB_2337972) or DTAF conjugated donkey anti-mouse (1:250, Jackson ImmunoResearch, Pennsylvania, United States, www.jacksonimmuno.com), signals could be directly observed under a fluorescence microscope (Eclipse E800M, Nikon). Whereas with secondary antibodies like biotinylated goat anti-rabbit (1:500, BA-1000, Vector Laboratories, California, United States, https://vectorlabs.com, RRID: AB_2313606) or biotinylated goat anti-mouse antibody (1:500, BA-9200, Vector Laboratories, California, United States, https://vectorlabs.com, RRID: AB_2336171), the signals were mostly amplified by the Avidin-Biotin-peroxidas complex (ABC kit, Vector) and further detected suing the Tyramide Signal Amplification (TSA, PerkinElmer) system. For antibody staining requires antigen retrieval, sections were heated at 95°C for 10 min in the citrate buffer (10 mM Citric Acid, pH 6.0), and cool down at room temperature before general IHC staining procedure.

For most of the double *in situ* hybridization/IHC experiments in this study, *in situ* was performed firstly and detected with the tyramide-FITC or tyramide-Cy3, whereas the later-performed IHC was detected using the tyramide-Cy3 or tyramide-FITC, respectively. However, the double labeling of *Zswim5* and LHX6 was performed using the NBT/BCIP system for the *in situ* hybridization of *Zswim5*, and the 3,3’-diaminobenzidine tetrahydrochloride (DAB, Sigma) system for the detection of LHX6. In brief, the *in situ* hybridization for *Zswim5* was performed first following the normal protocol. However, the probe used here was the full-length probe of *Zswim5* to ensure the staining of *Zswim5*-positive cells in the developing neocortex. For the IHC of LHX6, after the general amplification step of the ABC kit (0.5% in 0.1M PBS, Vector) for 1 hour, sections were washed in 0.1 M PBS. Then, the LHX6 signal is further amplified with the biotinylated (bio)-tyramide (BT) for 20 min and again with the ABC kit (0.15% in 1ml 0.1M PBS, Vector) for 1 hour. Finally, after washing with 0.1 M PBS, LHX6 is detected with the DAB system (Sigma). After the color development, the slides were treated with 50%, 70%, 90% and 100% acetone to bleach out the background mainly caused by the *in situ* hybridization. This procedure caused the color of *in situ* hybridization signals turned blue instead of the original dark purple.

## RESULTS

We performed *in situ* hybridization using digoxigenin-labeled riboprobes to characterize the developmental expression patterns of *Zswim5* transcripts from embryonic stages to adulthood of mouse telencephalon. In parallel, in most developmental stages, we also performed *in situ* hybridization using ^35^S isotope-labeled riboprobes to validate the expression pattern of *Zswim5* mRNA that was characterized with digoxigenin-labeled riboprobes. The specificity of *Zswim5* expression pattern was confirmed by performing the *in situ* hybridization using the sense riboprobes, which resulted in the non-specific background with no specific signals (data not shown).

Overall, *Zswim5* was expressed in the subventricular zone (SVZ) of the MGE, CGE, preoptic area (POA) and preoptic hypothalamic border domain (POH) during embryonic stages. *Zswim5* was subsequently down-regulated after birth. *Zswim5* was also moderately to weakly expressed in the cortical plate (CP), amygdala, thalamus, and hypothalamus in embryonic stages. Detailed expression patterns and expression levels of *Zswim5* mRNA are summarized in Table 2.

**Table 2.**
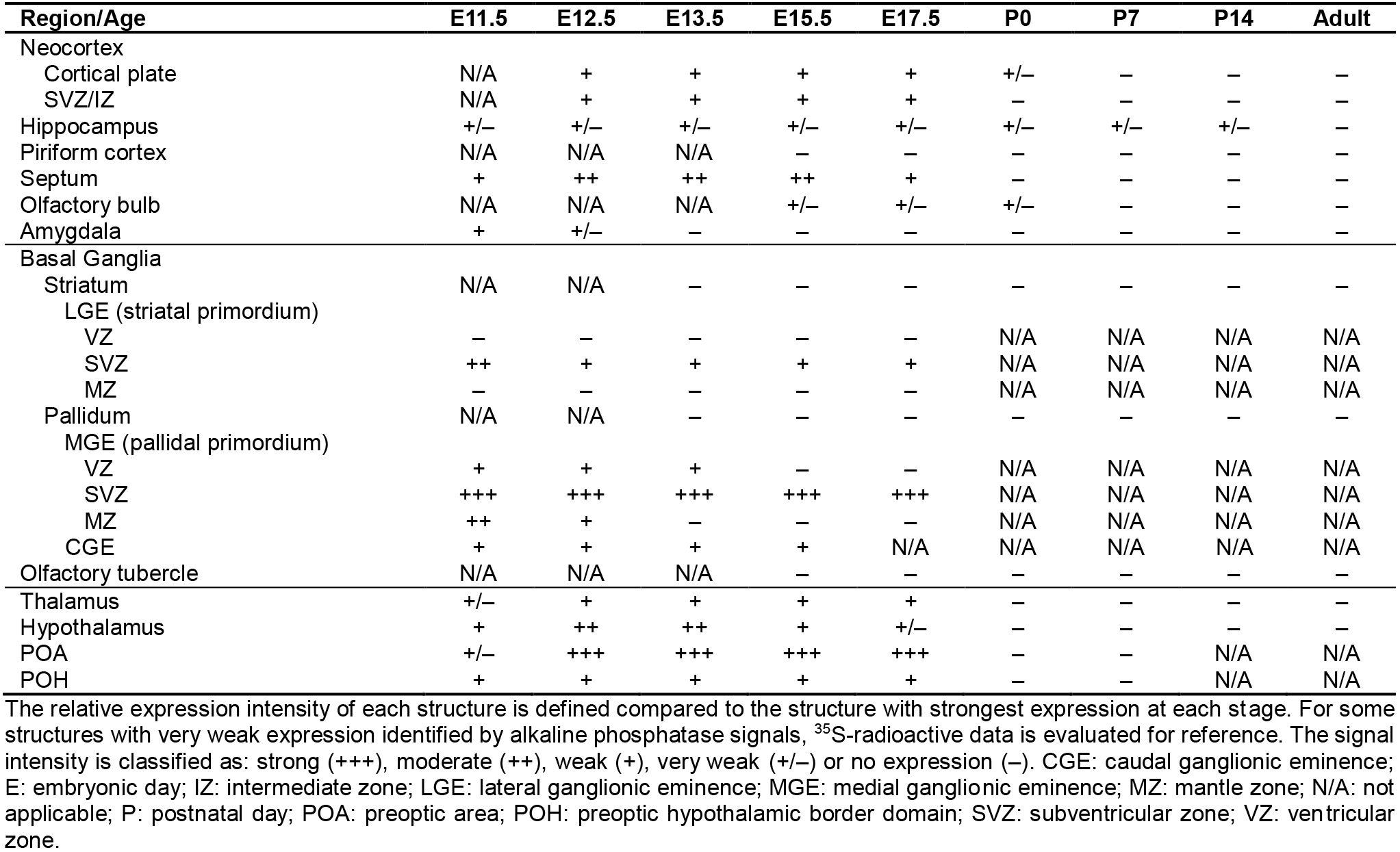
Expression patterns of Zswim5 in the developing mouse forebrain

| Region/Age | E11.5 | E12.5 | E13.5 | E15.5 | E17.5 | P0 | P7 | P14 | Adult |
| --- | --- | --- | --- | --- | --- | --- | --- | --- | --- |
| Neocortex |  |  |  |  |  |  |  |  |  |
| Cortical plate | N/A | + | + | + | + | +/- | - | - | - |
| SVZ/IZ | N/A | + | + | + | + | - | - | - | - |
| Hippocampus | +/- | +/- | +/- | +/- | +/- | +/- | +/- | +/- | - |
| Piriform cortex | N/A | N/A | N/A | - | - | - | - | - | - |
| Septum | + | ++ | ++ | ++ | + | - | - | - | - |
| Olfactory bulb | N/A | N/A | N/A | +/- | +/- | +/- | - | - | - |
| Amygdala | + | +/- | - | - | - | - | - | - | - |
| Basal Ganglia |  |  |  |  |  |  |  |  |  |
| Striatum | N/A | N/A | - | - | - | - | - | - | - |
| LGE (striatal primordium) |  |  |  |  |  |  |  |  |  |
| VZ | - | - | - | - | - | N/A | N/A | N/A | N/A |
| SVZ | ++ | + | + | + | + | N/A | N/A | N/A | N/A |
| MZ | - | - | - | - | - | N/A | N/A | N/A | N/A |
| Pallidum | N/A | N/A | - | - | - | - | - | - | - |
| MGE (pallidal primordium) |  |  |  |  |  |  |  |  |  |
| VZ | + | + | + | - | - | N/A | N/A | N/A | N/A |
| SVZ | +++ | +++ | +++ | +++ | +++ | N/A | N/A | N/A | N/A |
| MZ | ++ | + | - | - | - | N/A | N/A | N/A | N/A |
| CGE | + | + | + | + | N/A | N/A | N/A | N/A | N/A |
| Olfactory tubercle | N/A | N/A | N/A | - | - | - | - | - | - |
| Thalamus | +/- | + | + | + | + | - | - | - | - |
| Hypothalamus | + | ++ | ++ | + | +/- | - | - | - | - |
| POA | +/- | +++ | +++ | +++ | +++ | - | - | N/A | N/A |
| POH | + | + | + | + | + | - | - | N/A | N/A |

The relative expression intensity of each structure is defined compared to the structure with strongest expression at each stage. For some structures with very weak expression identified by alkaline phosphatase signals, <sup>35</sup>S-radioactive data is evaluated for reference. The signal intensity is classified as: strong (+++), moderate (++), weak (+), very weak (+/-) or no expression (-). CGE: caudal ganglionic eminence; E: embryonic day; IZ: intermediate zone; LGE: lateral ganglionic eminence; MGE: medial ganglionic eminence; MZ: mantle zone; N/A: not applicable; P: postnatal day; POA: preoptic area; POH: preoptic hypothalamic border domain; SVZ: subventricular zone; VZ: ventricular zone.

### *Zswim5* mRNA expression in embryonic mouse forebrain

#### E11.5

The expression pattern of *Zswim5* mRNA at E11.5 (n = 5) from rostral to caudal levels are illustrated in Figure 1 (A1-A6, B1-B10). At the rostral levels, *Zswim5* mRNA was first found to be weakly expressed in the septal area (Fig. 1A1, B1, B2, E), and extended up along the ventral telencephalon into the primordium of the striatum, which is known as the LGE (Fig. 1A1, B2). *Zswim5* expression in the LGE was mainly found in the SVZ (Fig.1A2, C, C1), which is the transition area while proliferating progenitor cells transform to differentiating cells and further entering the mantle zone (MZ). Weak signals of *Zswim5* were further extended up from the SVZ of LGE into the developing cortical primordium and then gradually decreased (Fig.1D1, D2).

**Figure 1.**
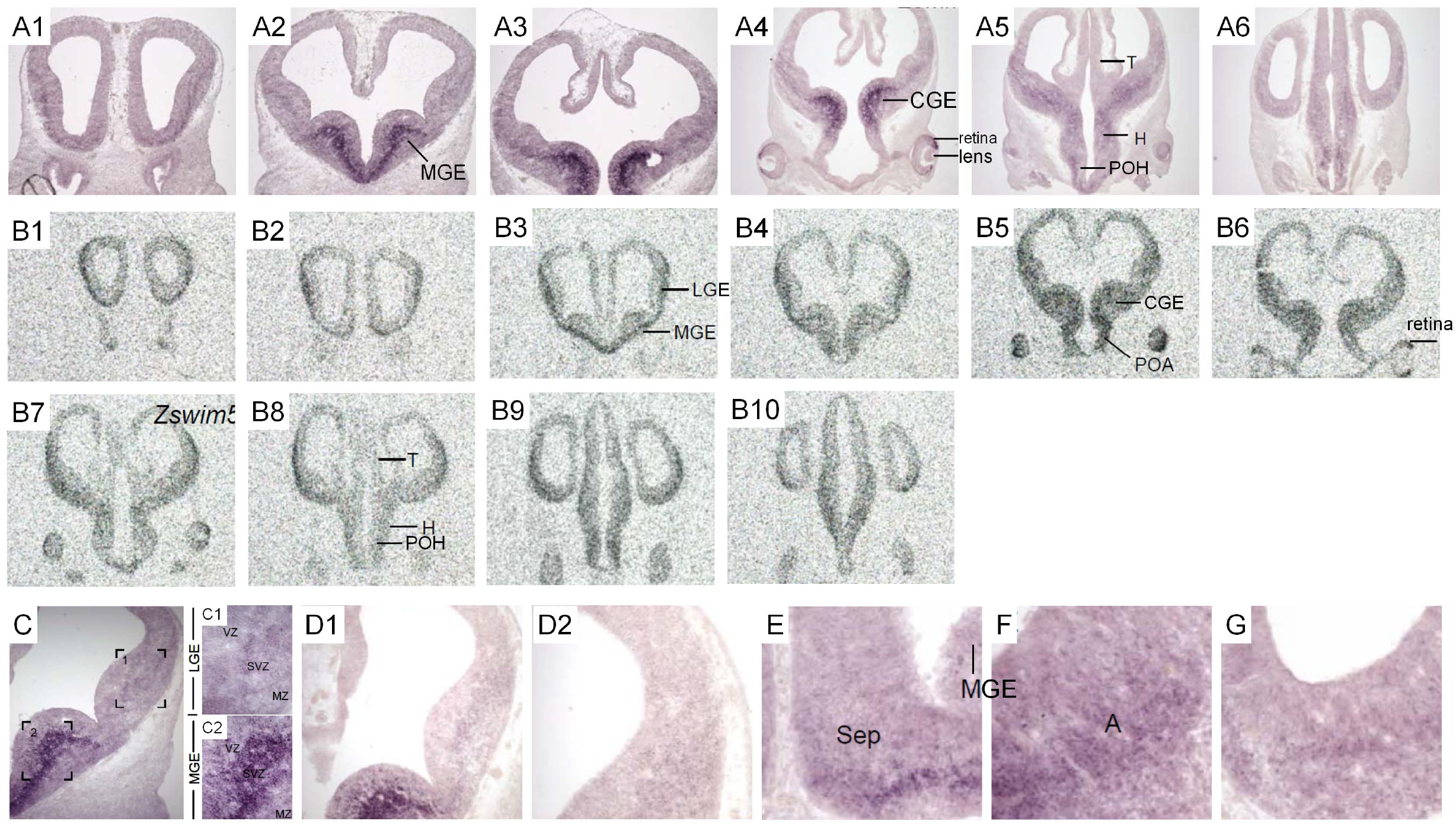
*Zswim5* mRNA expression pattern in E11.5 mouse forebrain. **(A, B)** Expression pattern of *Zswim5* mRNA assayed by *in situ* hybridization with digoxigenin-labeled probes (A1-A6) and ^35^S-labeled probes (B1-B10) from rostral to caudal levels. **(C, D)** *Zswim5* is strongly expressed in the SVZ of MGE but is weakly expressed in the SVZ of LGE. *Zswim5*-positive signals extend upward to the SVZ of LGE and gradually down-regulated at the border between LGE and cortical primordium at E11.5 (D1-2). **(E, G, F)** Expression pattern of *Zswim5* in the septum (E), amygdala (F) and entorhinal cortex (G). A: amygdala; CGE: caudal ganglionic eminence; H: hypothalamus; LGE: lateral ganglionic eminence; MGE: medial ganglionic eminence; MZ: mantel zone; POA: preoptic area; POH: preoptic hypothalamic border domain; Sep: septum; SVZ: subventricular zone; T: thalamus; VZ: ventricular zone.

In contrast to the weak expression of *Zswim5* in LGE, *Zswim5* was most strongly expressed in the pallidum primordium, also known as the MGE (Fig. 1A2-A4, Fig. 1C, C2). Comparing to many cells expressing strong *Zswim5* in the SVZ of MGE, a few scattered *Zswim5*-positive cells were found in the adjacent ventricular zone (VZ) (Fig. 1C2). Interestingly, *Zswim5* was also highly expressed in the POA as well as in the CGE. In fact, a continuous band of prominent *Zswim5* expression appeared to extend from the POA through the MGE and CGE into the overlying cortical primordium (Fig. 1A4, B5).

Structures ventral to the MGE, such as the POA and the eye primordium were also *Zswim5*-positive (Fig. 1A4, B5, B6). Strong *Zswim5* was detected in the front-most area of the retina, while no expression was found in the lens (Fig. 1A4, B6). In the caudal brain regions, *Zswim5* mRNA was detected in the POH, primordia of the amygdala, thalamus, hypothalamus, and the entorhinal cortex with weak expression levels (Fig. 1A5, A6, B8, F, G).

#### E12.5 and E13.5

The expression patterns of *Zswim5* at E12.5 (n = 10) and E13.5 (n = 7) from rostral to caudal levels are illustrated in Figure 2 (A1-A6, B1-B10) and Figure 3. The expression patterns of *Zswim5* at E12.5 and E13.5 were similar. At the rostral levels, *Zswim5* was moderately expressed in the septal primordium (Fig. 2A1, A2, B1-B3; Fig. 3A1, A2). The moderate signals from the septum were extended into the SVZ of LGE. At the level where the MGE started to appear, *Zswim5* expression was increased in the septum (Fig. 2A2; Fig. 3A3). In the MGE, scattered *Zswim5*-positive cells were present in the VZ of MGE (Fig. 2A2, A3, B4, B5; Fig. 3A3-A6). Strong *Zswim5* expression was found in the SVZ of MGE while the expression level was significantly decreased in the differentiating MZ (Fig. 2C, D1). Interestingly, a band of *Zswim5*-positive cells appeared to tangentially migrate from the SVZ of MGE, through the SVZ of LGE into the overlying cortical primordia (Fig. 2C1, D1). Within the cortical primordia, *Zswim5*-positive cells appeared to migrate along the subventricular zone/intermediate zone (SVZ/IZ) (Fig. 2D1, D2). The signal intensity of these *Zswim5*-positive cells entering the developing neocortex from the SVZ of LGE or CGE was not strong (Fig. 2D2). *Zswim5*-positive cells were also found in the cortical plate (CP) of the developing neocortex (Fig. 2D2). In addition, this *Zswim5*-positive stream of cells further entered the developing hippocampal primordium (Fig. 2F).

**Figure 2.**
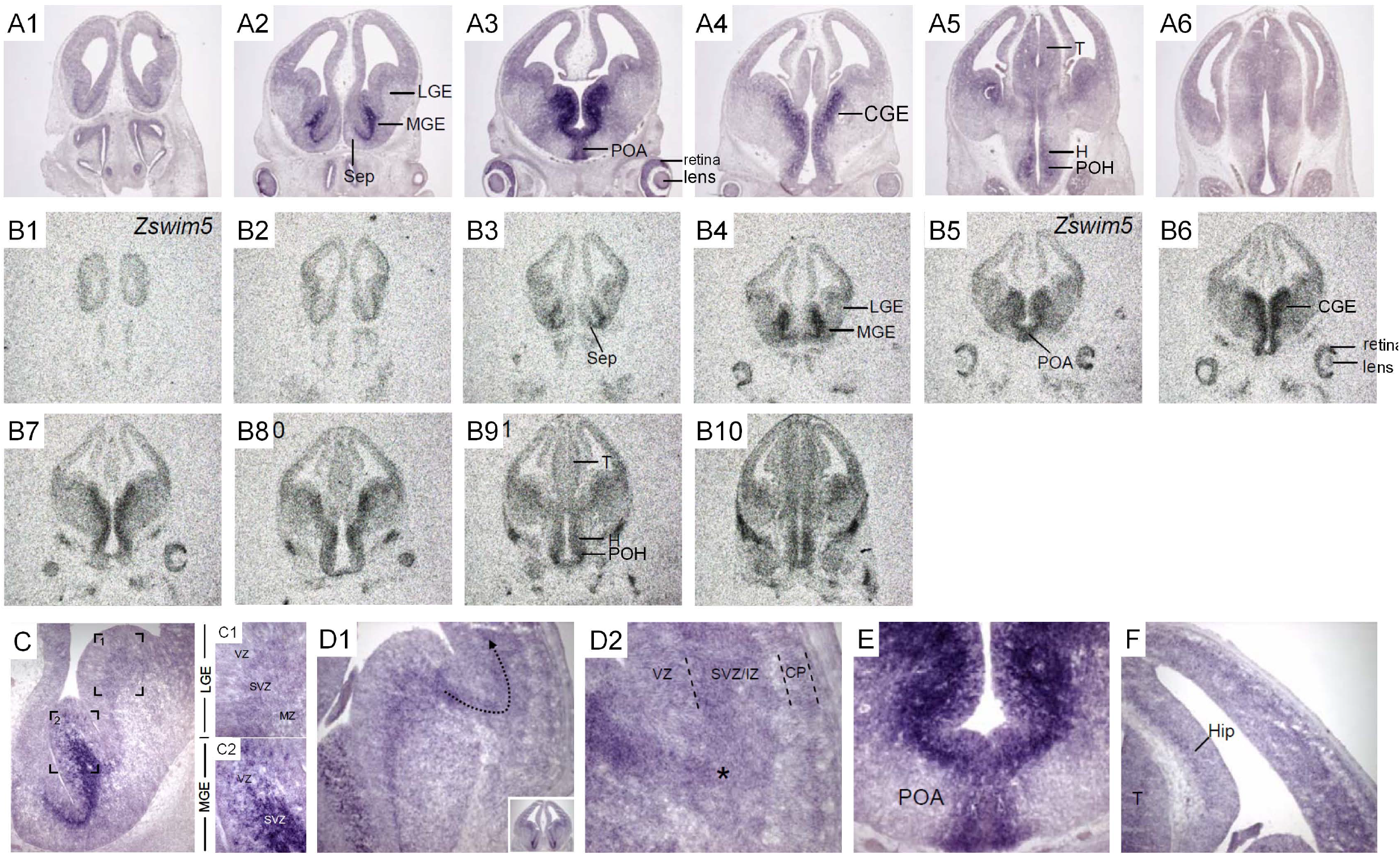
*Zswim5* mRNA expression pattern in E12.5 mouse forebrain. **(A, B)** Expression pattern of *Zswim5* mRNA assayed by *in situ* hybridization with digoxigenin-labeled probes (A1-A6) and ^35^S-labeled probes (B1-B10) from rostral to caudal levels. **(C, D)** *Zswim5* is highly expressed in the SVZ of MGE but is weakly expressed in the SVZ of LGE (C, C1, C2). *Zswim5* signals in the SVZ of MGE extend into the SVZ of LGE and then turn (star) along the corticostriatal sulcus to form tangentially arranged corridors of cells in the SVZ/IZ of the neocortex (D1, D2). A: amygdala; CGE: caudal ganglionic eminence; CP: cortical plate; H: hypothalamus; Hip: hippocampus; IZ: intermediate zone; LGE: lateral ganglionic eminence; MGE: medial ganglionic eminence; MZ: mantel zone; POA: preoptic area; POH: preoptic hypothalamic border domain; Sep: septum; SVZ: subventricular zone; T: thalamus; VZ: ventricular zone.

**Figure 3.**
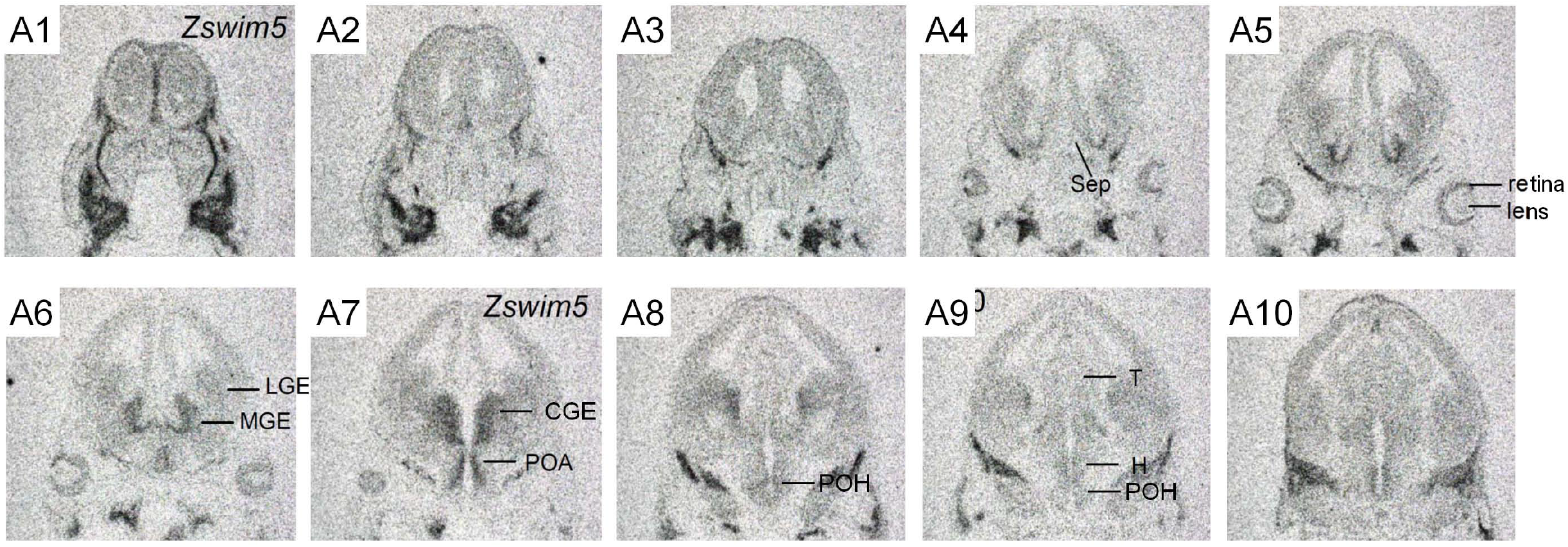
*Zswim5* mRNA expression pattern in E13.5 mouse forebrain. Expression pattern of *Zswim5* mRNA from rostral to caudal levels assayed by *in situ* hybridization with S^35^-labeled probes. CGE: caudal ganglionic eminence; H: hypothalamus; LGE: lateral ganglionic eminence; MGE: medial ganglionic eminence; POA: preoptic area; POH: preoptic hypothalamic border domain; T: thalamus.

Moderate levels of *Zswim5* mRNA was detected in the CGE (Fig. 2A4, 2B6, 3A7). Similar to that in the MGE, *Zswim5* was strongly expressed in the SVZ of POA and POH with a few *Zswim5*-positive cells in the VZ (Fig. 2A3, A5, B5, B9, E; Fig. 3A7, A8, A9). Strips of *Zswim5*-positive cells were found in the outermost part of the retina, while other cells in the retina had a homogeneous moderate expression of *Zswim5* (Fig. 2A3, B6; Fig. 3A5). Only a few cells in the lens were *Zswim5* positive (Fig. 2A3, B6). Weaker signals were found in the developing thalamus than that in the hypothalamus, which included the fields of Forel, ventral, intermediate and lateral hypothalamus (Fig. 2A5, A6, B8-B10; Fig. 3A9). Among these subdivisions of the hypothalamus, ventral hypothalamus had the highest expression.

#### E15.5

The expression patterns of *Zswim5* mRNA at E15.5 (n = 8) from rostral to caudal levels are illustrated in Figure 4 (A1-A7; B1-B9). *Zswim5* remained to be highly expressed in the SVZ of the pallidum primordium (Fig. 4A4, A5; B4). As in E12.5 and E13.5, the *Zswim5* signals in the pallidum primordium further extended up into the SVZ of the LGE and finally into the tangentially arranged corridors of cells in the SVZ/IZ of the neocortex (Fig. 4C1-C3). Cells in the head of the lateral migratory stream (Fig. 4C2, star) showed a prominent pathway of turning pattern along the corticostriatal boundary, supporting the finding that this stream of *Zswim5*-positive cells might be the tangentially migrating stream of cells, which migrate to the developing neocortex and form cortical interneurons. In addition, *Zswim5* was also evidently expressed in the CP, which becomes wider due to the packing of cortical neurons and interneurons migrating from neocortical neuroepithelium and various interneuron origins, respectively (Fig. 4C1, C3). These *Zswim5*-positive cells in the CP were also further extended into the medial cortical region and hippocampal primordium from rostral to caudal levels. Structures such as the striatum, thalamus, and hypothalamus contained low levels of *Zswim5*, whereas strong *Zswim5* expression remained in the POA (Fig. 4B6-B9). Low levels of *Zswim5* mRNA was detected in the CGE and POH (Fig. 4A6, B7).

**Figure 4.**
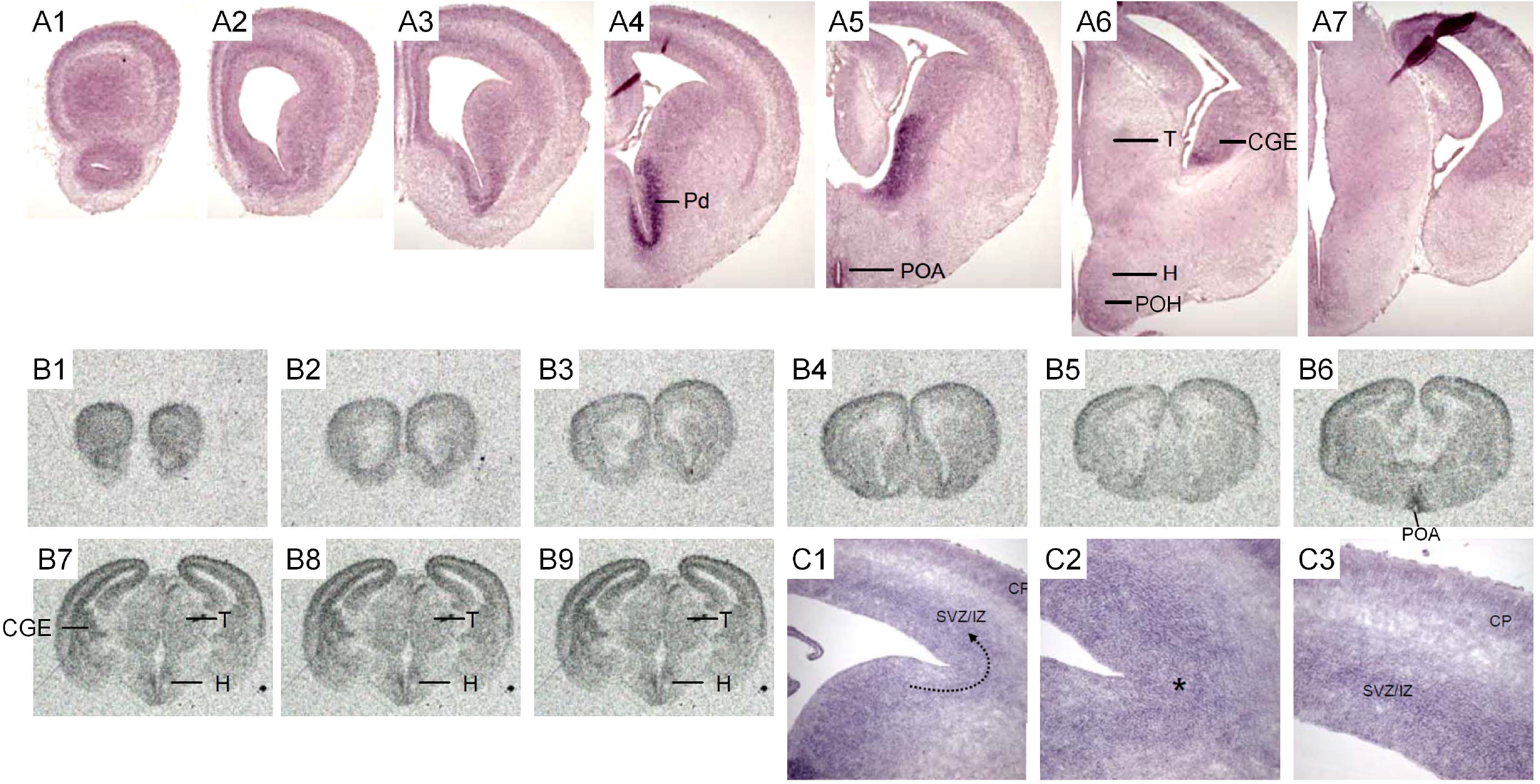
*Zswim5* mRNA expression pattern in E15.5 mouse forebrain. **(A, B)** Expression pattern of *Zswim5* mRNA assayed by *in situ* hybridization with digoxigenin-labeled probes (A1-A7) and S^35^-labeled probes (B1-B9) from rostral to caudal levels. **(C)** Similar to that at earlier stages, *Zswim5* signals in the SVZ of the residual pallidum primordium extend upward into the SVZ of LGE and then turn (star) along the corticostriatal sulcus to form tangentially arranged corridors of cells in the SVZ/IZ of the neocortex. A: amygdala; CGE: caudal ganglionic eminence; CP: cortical plate; H: hypothalamus; IZ: intermediate zone; IZ: intermediate zone; MGE: medial ganglionic eminence; Pd: pallidum primordium; POA: preoptic area; POH: preoptic hypothalamic border domain; VZ: subventricular zone; T: thalamus.

#### E17.5

The expression pattern of *Zswim5* mRNA at E17.5 (n = 5) from rostral to caudal levels are illustrated in Figure 5 (A1-A6; B1-B9). At rostral levels, *Zswim5* mRNA was not detected in the primordium of OB, (Fig. 5A1, B1). *Zswim5* remained strongly expressed in the SVZ of the residual pallidum primordium, which might represent the SVZ of nucleus accumbens (Acb) at late embryonic stages (Fig. 5A4). As in previous stages, *Zswim5* signals were further extended up into the SVZ/VZ of the developing striatum with a moderate level, passed through the corticostriatal boundary (Fig. 5C2, star) and entered the SVZ/IZ and VZ of the developing neocortex (Fig. 5C1-C3). As the developmental stages progressed, the SVZ/IZ and VZ of the cortical primordium gradually became thinner. As a result, the band of *Zswim5*-positive cells expressed in the SVZ of cortical primordium became thinner than that in E15.5 (Fig. 5C1, C3). In addition, *Zswim5* was expressed weakly along with the cortical plate ventrally to the piriform cortex and olfactory tubercle, and dorsomedially entered the cingulate cortex, which had slightly higher expression intensity (Fig. 5A2-A4; Fig. 5B3). *Zswim5* in the developing striatum showed very weak and disperse expression (Fig. 5A2-A4; B2-B5). At caudal levels, where the cortical plate extended medially into the hippocampus primordium, *Zswim5* signal was weakly expressed in the pyramidal cell of the hippocampus and in granular cells of the dentate gyrus (Fig. 5A5, A6; B5-B8). Other structures such as the thalamus, hypothalamus and amygdala were largely *Zswim5*-negative (Fig. 5A5, A6, B6, B7).

**Figure 5.**
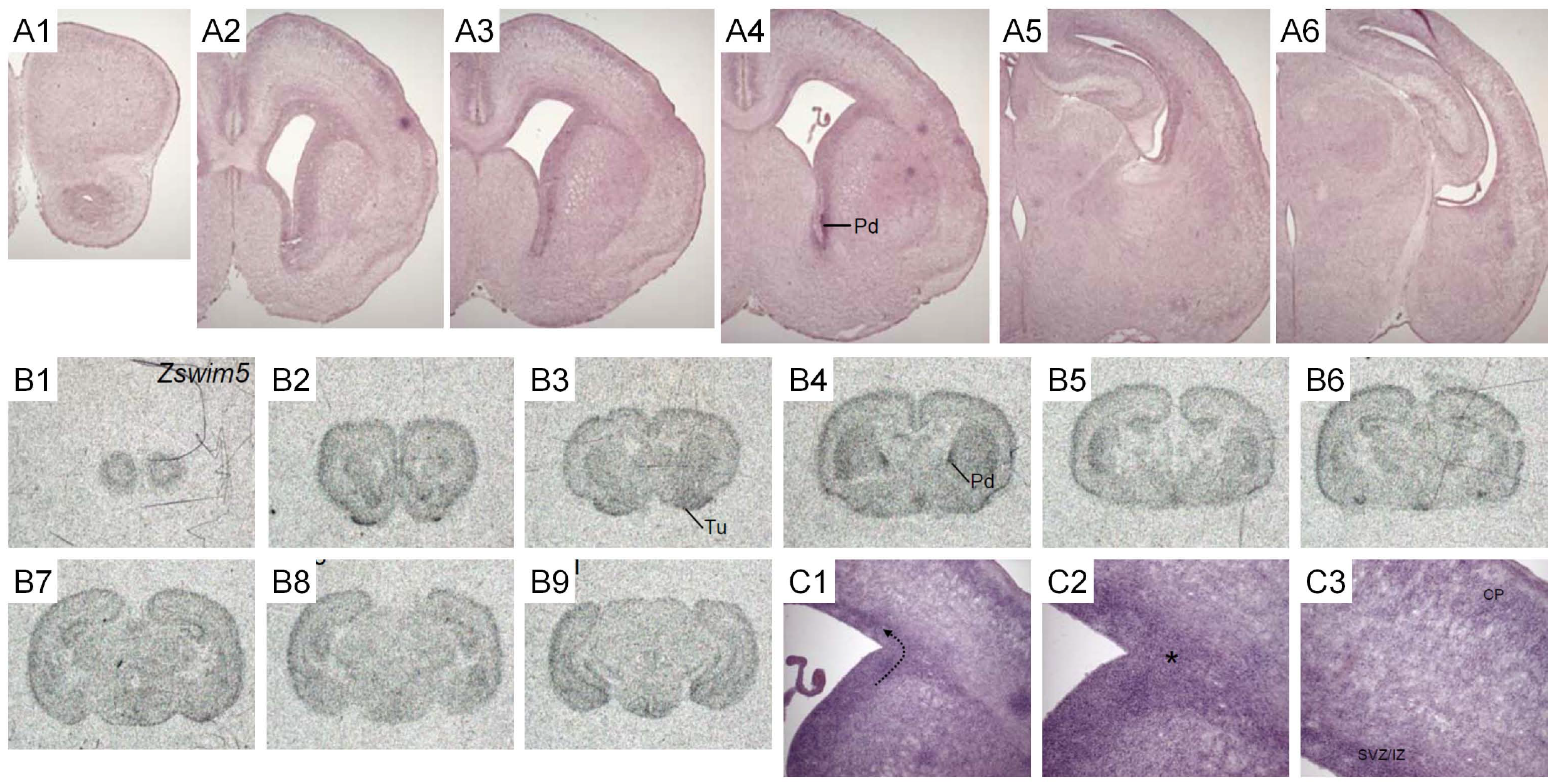
*Zswim5* mRNA expression pattern in E17.5 mouse forebrain. **(A, B)** Expression pattern of *Zswim5* mRNA assayed by *in situ* hybridization with digoxigenin-labeled probes (A1-A7) and ^35^S-labeled probes (B1-B9) from rostral to caudal levels. In the subpallium, *Zswim5* expression is decreased in the residual pallidum primordium at E17.5. In the developing neocortex, *Zswim5* is expressed at high levels in the SVZ/IZ. **(C)** Similar to the pattern at earlier stages, *Zswim5* signals in the SVZ of the residual pallidum primordium appear to extend into the SVZ of LGE and then turn (star) along the corticostriatal sulcus to enter the SVZ/IZ of the neocortex. IZ: intermediate zone; CP: cortical plate; Pd: pallidum primordium; SVZ: subventricular zone; T: thalamus; Tu: olfactory tubercle.

### *Zswim5* mRNA expression in postnatal mouse forebrain

The expression pattern of *Zswim5* mRNA at P0 (n = 3) from rostral to caudal levels are illustrated in Figure 6. Weak *Zswim5* expression remained in the SVZ of the residual pallidum primordium (or the SVZ of the Acb) (Fig. 6A3, A4). In general, the overall signal intensity of *Zswim5* at P0 was much lower than that at earlier stages. *Zswim5* expression was detected in the OB. Using more sensitive ^35^S-labeled riboprobes, low levels of *Zswim5* were further detected in the olfactory tubercle, the striatum, and hippocampus (Fig. 7B1-B4).

**Figure 6.**
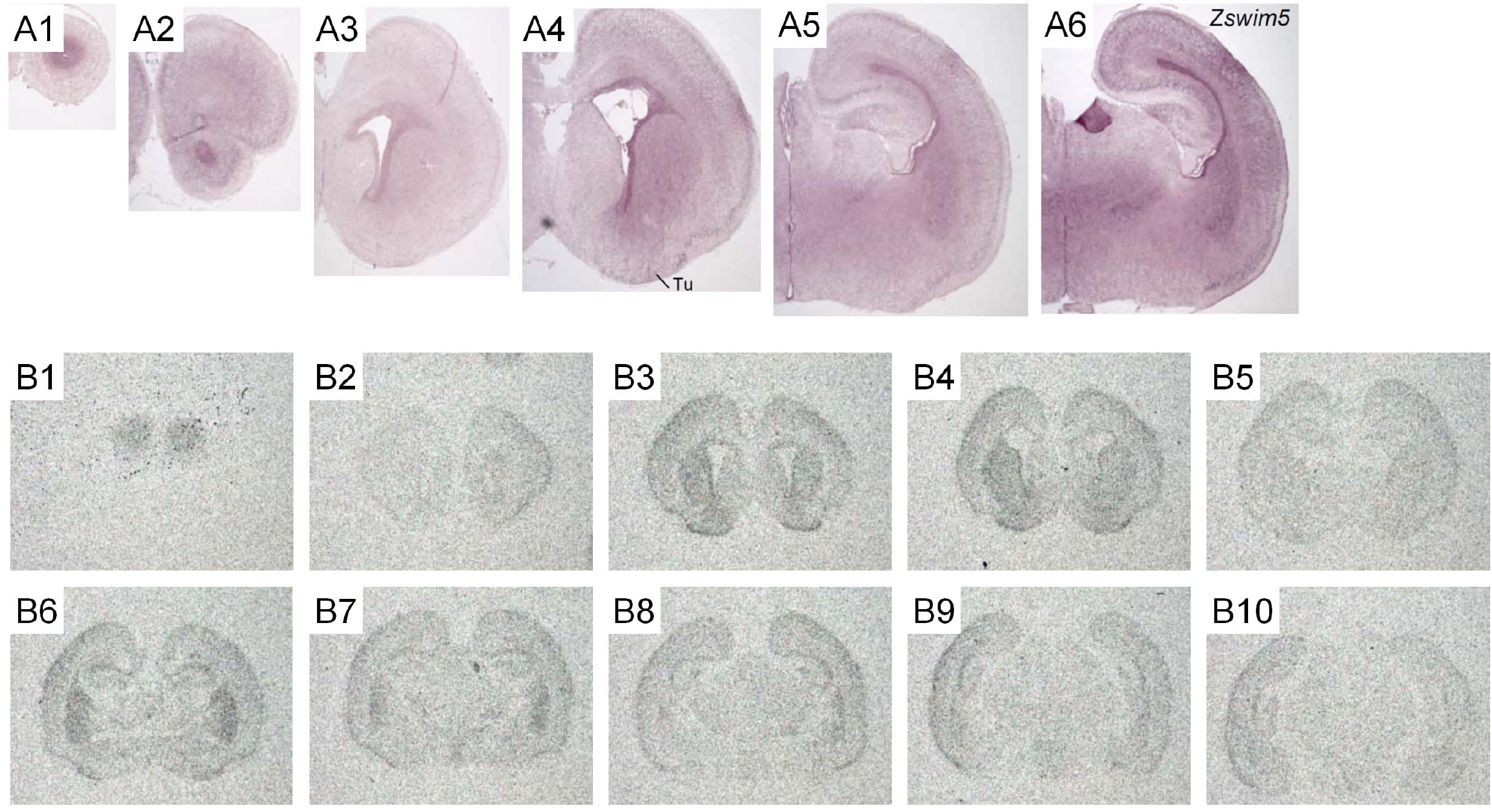
*Zswim5* mRNA expression pattern in P0 mouse forebrain. Expression pattern of *Zswim5* mRNA assayed by *in situ* hybridization with digoxigenin-labeled probes (A1-A6) and ^35^S-labeled probes (B1-B10) from rostral to caudal levels. Moderate *Zswim5* signals are detected in the olfactory bulb, cortex, and striatum.

**Figure 7.**
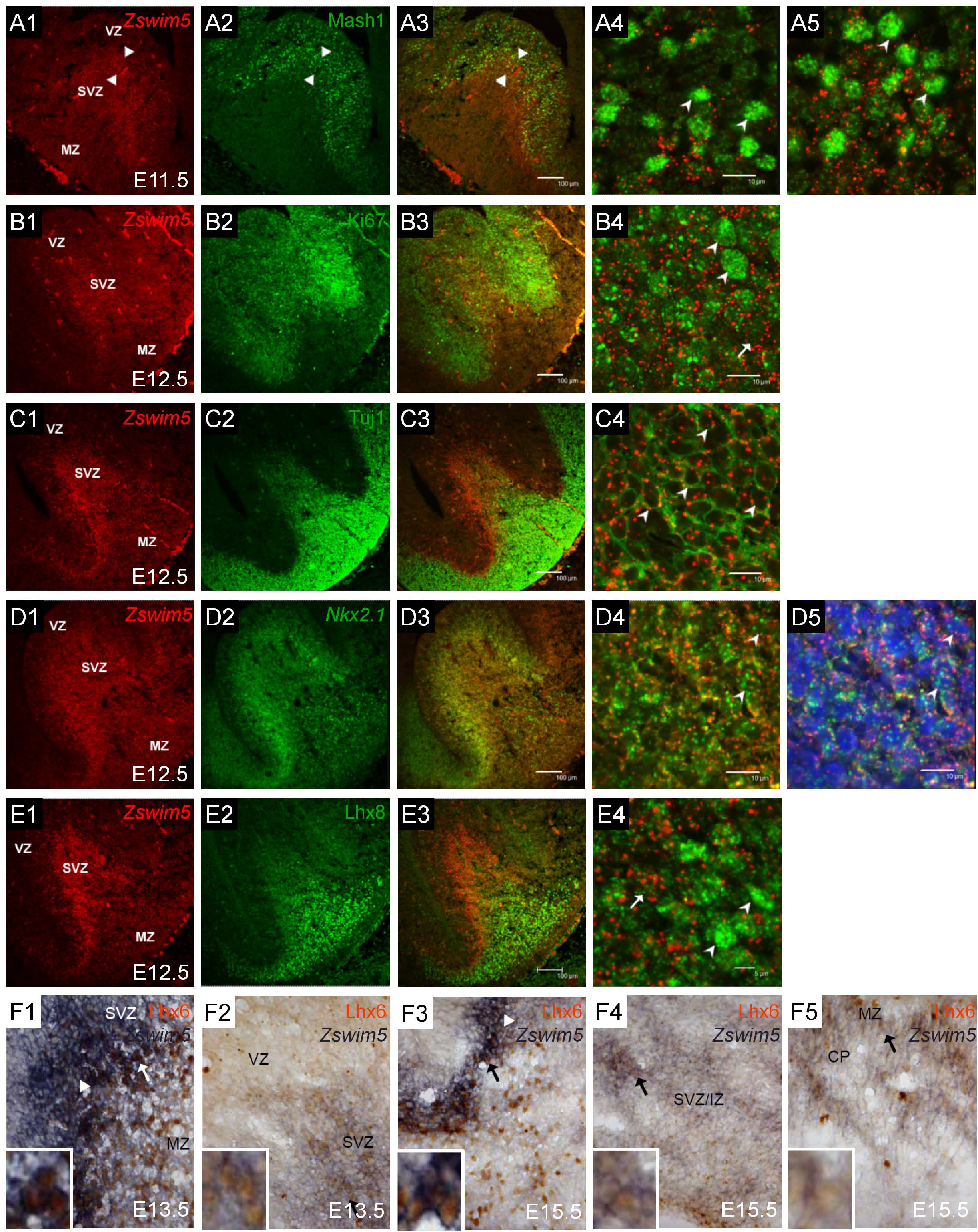
*Zswim5* is expressed by Lhx6-positive post-mitotic neurons that are originated from Nkx2.1-positive MGE. *Zswim5* is mainly expressed in the SVZ of MGE at E11.5 (A1) and E12.5 (B1, C1, D1, E1), whereas Mash1-positive progenitors (A2) and Ki67-positive proliferating neurons (B2) are mainly located in the VZ and the SVZ of MGE. In the overlapped zone, Mash1-positive (A3-A5, arrowheads) or Ki67-positive cells (B3-B5, arrowheads) are found to co-express none or at most low levels of *Zswim5*. In contrast, differentiating neurons expressing Tuj1 are located in the SVZ and MZ of MGE (C2, C3). Tuj1 immunoreactivity is detected in the cytoplasm (C4), and *Zswim5* mRNA are also detected in the cytoplasm as red granule puncta surrounding the nucleus (C4). Nkx2.1-positive cells are mainly located in the VZ and SVZ of MGE (D2, D3), and *Zswim5*-positive cells co-expressing Nkx2.1 mRNA are present in the SVZ of MGE (D2-D5, arrowheads). Lhx8 expressing cells (E2, E3) are detected in the SVZ and MZ of MGE, but Lhx8-positive cells show none (E4, arrowheads) or little *Zswim5* mRNA (E4, arrow) at E12.5. Lhx6 expressing GABAergic interneurons (F1-F5, brown) are mainly located in the SVZ and MZ of the MGE (F1; E13.5), LGE (F2; E13.5), developing striatum (F3; E15.5) and developing neocortex (F4, F5; E15.5). Co-localization of *Zswim5* mRNA (dark purple) and Lhx6 immunoreactivity are found in the SVZ of MGE (F1; strong: arrow; weak: arrowhead; magnification: inset), LGE (F2, inset), and pallidum primordium (F3, inset). *Zswim5* and Lhx6 double-positive cells are found in the SVZ/IZ (F4, inset) and the MZ/CP (F5, inset) of the developing neocortex, which presumably represents the population of tangential migratory neurons that eventually develop into interneurons in the cortex. Insets in F1-F5 show the magnifications of the cells indicated by arrows. CP: cortical plate; IZ: intermediate zone; MZ: mantle zone; SVZ: subventricular zone; VZ: ventricular zone. Scale bars: 100 μm in A3, B3, C3, D3, E3; 10 μm in A4, B4, C4, D4, D5; 5 μm in E4.

*Zswim5* signals were not detectable in the P7, P14 and adult sections except little *Zswim5* expression was found in the hippocampus (data not shown).

### *Zswim5* was expressed in early postmitotic neurons at embryonic stages

To examine if *Zswim5* was expressed in the population of proliferating progenitors, *Zswim5* mRNA was double labeled with the proneural gene *Mash1/Ascl1* and the proliferating marker Ki67 in E11.5 and E12.5 mouse brain, respectively. *Zswim5* mRNA was expressed homogeneously in the SVZ of MGE (Fig. 7A1), whereas Mash1 was primarily expressed in the VZ with low levels in the SVZ (Fig. 7A2). As a consequence, *Zswim5* and *Mash1* expressions were partially overlapped at the boundary between the VZ and the SVZ of MGE (Fig. 7A1-A3, between arrowheads). At the single-cell level, however, Mash1-positive cells appeared not to express *Zswim5* (Fig. 7A4-A5). Consistently, *Zswim5* expressing cells showed none or at most low levels of Ki67, though *Zswim5* and Ki67 shared an overlapping expression zone in the SVZ of MGE (Fig. 7B1-B4). These results suggested that *Zswim5* was likely to be immediately up-regulated upon the progenitors exiting the cell cycle in the MGE.

Tuj1, also known as the neuron-specific class III β-tubulin, is a marker for early differentiating neurons (Menezes and Luskin, 1994). To examine if *Zswim5* was expressed by differentiating progenitor, *Zswim5* mRNA was double labeled with Tuj1 at E12.5. *Zswim5* was expressed in the SVZ of MGE (Fig. 7C1), whereas Tuj1 was expressed in differentiating cells throughout the SVZ and the MZ of MGE (Fig. 7C2). At the single-cell level, it was evident that many *Zswim5*-positive cells co-expressed Tuj1 (Fig. 7C3, 7C4, arrowheads). Taken together, these results indicated that *Zswim5* was expressed in the postmitotic progenitors at early stages of neuronal differentiation.

### *Zswim5* was co-expressed by Nkx2.1 and Lhx6-positive neurons

Regarding *Zswim5* expression in MGE progenitors, *Zswim5* expression was detected in several MGE neuronal types. *Nkx2.1* not only plays a vital role in the formation of MGE (Sussel et al., 1999) but also is involved in the specification of two major populations of cortical GABAergic interneurons originating from MGE, including parvalbumin and somatostatin interneurons (Du et al., 2008). *Zswim5* mRNA and *Nkx2.1* mRNA expressions were overlapped in the SVZ of MGE (Fig. 7D1-D3), and *Zswim5*-positive cells co-expressing *Nkx2.1* were found at the single-cell level (Fig. 7D4, D5, arrowheads). Therefore, *Zswim5* expressing cells represented a subpopulation of *Nkx2.1*-positive progenitors that were located in the SVZ of MGE during development.

*Lhx6* is a direct downstream gene of *Nkx2.1* to specify cortical GABAergic interneurons that contributes to the proper migration of these interneurons throughout tangential migratory streams (Liodis et al., 2007; Du et al., 2008). To examine whether *Zswim5* was expressed in the progenitor population of cortical interneurons during development, *Zswim5* mRNA was immunostained with Lhx6 at E13.5 and E15.5. Some *Zswim5*-positive cells in the SVZ of MGE co-expressed with Lhx6 (Fig. 7F1, inset). In particular, *Zswim5*-positive cells located near the MZ had stronger Lhx6 expression (Fig. 7F1, arrow, inset), whereas *Zswim5*-positive cells close to the VZ showed weaker Lhx6 expression (Fig. 7F1, arrowhead). In the MZ of MGE, where *Zswim5* was not expressed, cells with strong Lhx6 expression were still evident to find (Fig. 7F1). In the SVZ of LGE, some *Zswim5*-positive cells also showed weak Lhx6 staining (Fig. 7F2, inset); whereas in the MZ of LGE, *Lhx6*-positive cells expressed a high level of Lhx6 but with weak or no *Zswim5* expression. At E13.5, a few *Lhx6*-positive or *Zswim5*-positive cells were found in the developing cortical primordium. The identification of *Zswim5/Lhx6* co-expressing cells was difficult in the developing cortex due to weak signals. By E15.5, *Zswim5* remained to be expressed in the SVZ of the residual pallidum primordium (Fig. 7F3, blue), and some *Zswim5*-positive cells co-expressed Lhx6 at high (Fig. 7F3, arrow, inset) or low levels (Fig. 7F3, arrowhead). Besides in the residual of pallidum primordium, *Zswim5*-positive cells in the MZ, CP or SVZ/IZ of the developing neocortex also co-expressed Lhx6 at low levels (Fig. 7F4, F5, insets). Taken together, these findings indicated that *Zswim5*-positive cells contribute to a subpopulation of the *Lhx6*-positive progenitors, which generated cortical GABAergic interneurons.

*Lhx8* is a LIM-homeobox transcription factor known to be specifically expressed in the MGE, the MGE-derived basal forebrain, and oral mesenchyme (Manabe et al., 2005). *Lhx8* plays a pivotal role in the development and maintenance of cholinergic neurons in the basal forebrain (Mori et al., 2004). *Zswim5* mRNA was mainly expressed at high levels in the SVZ of MGE (Fig. 7E1), whereas Lhx8 was detected primarily in the differentiated MZ of MGE (Fig. 7E2). Despite the partial overlap in the SVZ/MZ boundary (Fig. 7E1-E3), *Zswim5* mRNA and Lhx8 were not co-localized as examined at the single-cell level (Fig. 7E3, 7E4, arrowheads).

## DISCUSSION

This is the first study to comprehensively characterize the expression pattern of *Zswim5* mRNA in the developing mouse forebrain. In the early stages of forebrain development from E11.5 to E13.5, high level of *Zswim5* mRNA was primarily detected in the SVZ of the MGE and POA of the ventral forebrain. Based on the results of double labeling of Zswim 5 and proliferating or early differentiating markers, *Zswim5* is likely to be immediately upregulated as the progenitors exiting the cell cycle at the transition between proliferation and postmitotic differentiation. At E15.5 and E17.5, prominent expression of *Zswim5* remained detectable in the SVZ of the pallidal primordium (MGE). From neonatal to adult stages, *Zswim5* expression was drastically decreased in the forebrain.

The major finding of our study is that the expression pattern of *Zswim5* resembles the routes of tangential migration pathways. That is, progenitor cells in the MGE migrate through the SVZ of LGE and enter the SVZ/IZ of the developing neocortex to become cortical GABAergic interneurons. Interestingly, Nkx2.1 and *Zswim5* share a highly similar expression pattern in the MGE and POA (Sussel et al., 1999). We further found that *Zswim5* was co-localized with Nkx2.1 and Lhx6 in the MGE. As Nkx2.1 and Lhx6 expressed mainly in the MGE are noted for their roles in regulating the tangential migration and the specification of PV and SST interneurons (Du et al., 2008), our findings raise an interesting possibility that *Zswim5* may be a downstream target gene of *Nkx2.1*. *Nkx2.1* is one of the earliest regulatory genes that are at the upstream of the genetic cascades in the developmental regulation of forebrain neurons. The *Nkx2.1* null mutation causes failure in the formation of the MGE derivative due to a ventral to dorsal molecular re-specification in the MGE, and further causes the loss of GABAergic and calbindin-positive interneurons in the cortex and cholinergic interneurons in the striatum (Sussel et al., 1999). Moreover, *Nkx2.1* specifies the fate of cortical interneurons through direct activation of *Lhx6*, which is also responsible for regulating normal migration of the MGE-derived progenitor cells into the developing cortex (Liodis et al., 2007; Du et al., 2008). Thus, our findings that *Zswim5* was co-localized with both Nkx2.1 and Lhx6 suggest a role of *Zswim5* participating in the tangential migrating mechanism.

*Zswim5* appears to be expressed in the tangential migration pathways of Nkx2.1 lineages. Nkx2.1 is expressed in proliferating and postmitotic progenitors in the VZ and SVZ of MGE, and Lhx6 is expressed in cells after their last cell division in the SVZ and mantel zone of the MGE (Lavdas et al., 1999; Liodis et al., 2007). Given that *Zswim5* is mainly expressed in the early differentiating progenitors in the SVZ of MGE, it is possible that Nkx2.1 may activate both *Zswim5* and Lhx6. The *Zswim5*+/Lhx6+ progenitor cells may follow the tangential migrating routes via SVZ/IZ into the developing cortex. The specificity of *Zswim5*-positive progenitors for developing cortical interneurons is further supported by the finding that *Zswim5* was not expressed in Lhx8-positive progenitors that develop into cholinergic neurons in the basal forebrain (Manabe et al., 2005).

Our findings indicate that *Zswim5* is expressed in the progenitor domains of cortical GABAergic interneurons, including the MGE, CGE, POA and POH. Previous studies have reported that distinct domains contain the progenitors of different subtypes of cortical interneurons, i.e., The MGE and POA contains the progenitors of PV- and SST-positive interneurons, whereas the CGE and POH comprises the progenitors of RELN-positive, VIP-positive and other 5HT3a receptor-positive interneurons (Miyoshi et al., 2013; Marín, 2015; Bandler et al., 2017; Hu et al., 2017; Lim et al., 2018). Given that *Zswim5* is expressed throughout the MGE, CGE, POA and POH, *Zswim5* may be involved in the regulation of cortical interneurons derived from these regions, including PV-, SST- and 5HT3a receptor-positive interneurons. Future study of the genetic fate mapping of *Zswim5*-positive lineages and functional studies should help clarify the role of *Zswim5* in the regulation of development of cortical GABAergic interneurons.

Recent studies have characterized the cell lineages of cortical interneurons using the high-throughput technology of single-cell RNA sequencing, and *Zswim5* has been identified as a progenitor marker of cortical interneurons (Mayer et al., 2018; Mi et al., 2018). These single-cell RNA-seq studies show that *Zswim5* is expressed at a higher level in progenitors than that in neurons with computational analyses. Our current histological study confirmed that *Zswim5* is indeed expressed in the progenitor domains of cortical interneurons at early stages of development. The double *in situ* hybridization and immunostaining experiment further demonstrated that *Zswim5* was not co-localized in strong Ki67-positive proliferating progenitors in the SVZ, but was co-expressed in cells containing none or at most low levels of Ki67 (Fig. 7B1-B4) In contrast, *Zswim5* was colocalized in differentiating Tuj1-positive neurons in the SVZ (Fig. 7C1-C4). These results suggest that *Zswim5* is upregulated in cortical interneuron progenitors that are at the transition from exiting the cell cycle to postmitotic differentiation. Mayer et al. (2018) have reported that transcriptional profiles are largely conserved with a moderate difference across the three ganglionic eminences in progenitors of interneurons, the initial diversity of immature postmitotic neurons has already determined their cell type identity (Mayer et al., 2018). Consistently, Mi et al. (2018) have shown that the cell fate of cortical interneurons are intrinsically determined shortly after become postmitotic before they reach their final destinations (Mi et al., 2018). Along this line, the timing of the initiation of *Zswim5* expression in early postmitotic neurons is of particular interest, which suggests that *Zswim5* may play a role in cell fate determination of cortical interneurons.

In summary, our study has characterized the spatial and temporal expression pattern of *Zswim5* transcript in the developing mouse forebrain. Considering that *Zswim5* is upregulated in early differentiating cortical GABAergic interneurons, *Zswim5* may be potentially involved in the regulation of cell fate determination, migration and differentiation. Future studies using conditionally genetic manipulation of *Zswim5* may uncover the biological function of *Zswim5* in the developmental regulation of cortical interneurons.

## ACKNOWLEDGEMENTS

We thank Drs. H. Koga and T. Nagase for providing the *Zswim5* cDNA clone (mKIAA1511). This work was supported by the Ministry of Science and Technology-Taiwan grants MOST107-2321-B-010-002, MOST107-2320-B-010-041-MY3, the Featured Areas Research Center Program within the framework of the Higher Education Sprout Project by the Ministry of Education in Taiwan (F.-C.L.), and Postdoctoral Fellowship grants MOST107-2811-B-010-011, MOST107-2321-B-010-010-MY3 (H.-Y.K.).

## DATA AVAILABILITY STATEMENT

Data sharing is not applicable to this article as no new database was created or analyzed in this study.

